# Genetic variability in the expression of the SARS-CoV-2 host cell entry factors across populations

**DOI:** 10.1101/2020.04.06.027698

**Authors:** Lourdes Ortiz-Fernández, Amr H Sawalha

**Author notes:** Please address correspondence to Amr H. Sawalha, MD. Address: 7123 Rangos Research Center, 4401 Penn Avenue, Pittsburgh, PA 15224, USA.

## Abstract

The entry of SARS-CoV-2 into host cells is dependent upon angiotensin-converting enzyme 2 (ACE2), which serves as a functional attachment receptor for the viral spike glycoprotein, and the serine protease TMPRSS2 which allows fusion of the viral and host cell membranes. We devised a quantitative measure to estimate genetic determinants of *ACE2* and *TMPRSS2* expression and applied this measure to >2,500 individuals. Our data show significant variability in genetic determinants of *ACE2* and *TMPRSS2* expression among individuals and between populations, and demonstrate a genetic predisposition for lower expression levels of both key viral entry genes in African populations. These data suggest that genetic factors might lead to lower susceptibility for SARS-CoV-2 infection in African populations and that host genetics might help explain inter-individual variability in disease susceptibility and severity of COVID-19.

## Main Text

The severe acute respiratory syndrome coronavirus 2 (SARS-CoV-2) is a novel single-stranded RNA virus of the *Coronaviridae* family. This recently emerged virus is the cause of a pandemic infection that can result is severe life-threatening disease (Coronavirus Disease-2019; COVID-19).^1,2^ Similar to SARS-CoV (which causes SARS), SARS-CoV-2 entry into target host cells is mediated through binding of the viral spike glycoprotein to angiotensin-converting enzyme 2 (ACE2) on the cell surface.^3^ The interaction between coronaviral spike proteins and their attachment receptor ACE2 is believed to be key for viral transmissibility and dissemination of infection to organs and tissues.^4^ In addition, the host cell serine protease TMPRSS2 cleaves the spike protein of SARS-CoV-2 to allow fusion of the viral and host cell membranes, which is an essential step in viral entry.^3^

To gain insight into genetic determinants of SARS-CoV-2 transmissibility and potential viremia and disseminated infection, we devised a quantitative measure to assess the cumulative effect of genetic variants upon the expression of *ACE2* and *TMPRSS2*, and then applied this measure to determine differences in >2,500 individuals from 5 different populations around the world.

We used expression quantitative trait loci (eQTL) data for *ACE2* and *TMPRSS2* in tissues included in the Genotype-Tissue Expression project (GTEx, release V8),^5^ and derived and compared a genetic risk expression score (GRES) across all populations included in the 1000 Genomes Project.^6^ All eQTL variants affecting *ACE2* and *TMPRSS2* expression across all cell types and tissues were identified, and then pruned to remove variants in linkage disequilibrium (LD). LD pruning was performed using PLINK v.1.9 and the combined haplotypes of the 1000 Genomes Project populations.^7^ A total of 21 genetic polymorphisms that affect *ACE2* expression, and 14 that affect *TMPRSS2* expression were identified and used in subsequent analysis. The effect of each polymorphism upon corresponding gene expression was determined using an additive model for each gene separately and derived by multiplying the number of alternative alleles in each individual by the corresponding eQTL normalized effect size for each variant. For variants that demonstrate eQTL effects in multiple tissues, the most significant eQTL normalized effect size value was used. A cumulative genetic risk expression score was derived using the formula: 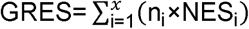, where n is the number of alternative alleles (0, 1 or 2), x is the number of evaluated variants in *ACE2* and *TMPRSS2* (21 and 14, respectively), and NES is the normalized effect size which reflects the expression in the alternative allele relative to the reference allele for each variant. The genetic variants used to calculate GRES values for *ACE2* and *TMPRSS2* are shown in **Supplementary Table 1**.

We compared GRES values for *ACE2* and *TMPRSS2* in 2,504 individuals across the 5 major populations included in the 1000 Genomes project (African, American, European, East Asian, and South Asian)^6^. Because *ACE2* is located on the X chromosome, we evaluated female and male individuals separately. There was a significant difference in the genetic risk expression score of *ACE2* between populations (ANOVA *P* <0.0001 for both male and female groups). Genetic determinants of highest expression of *ACE2* were observed in South Asian and East Asian populations, while African populations are genetically associated with the lowest *ACE2* expression levels (**Figure 1**).

**Figure 1:**
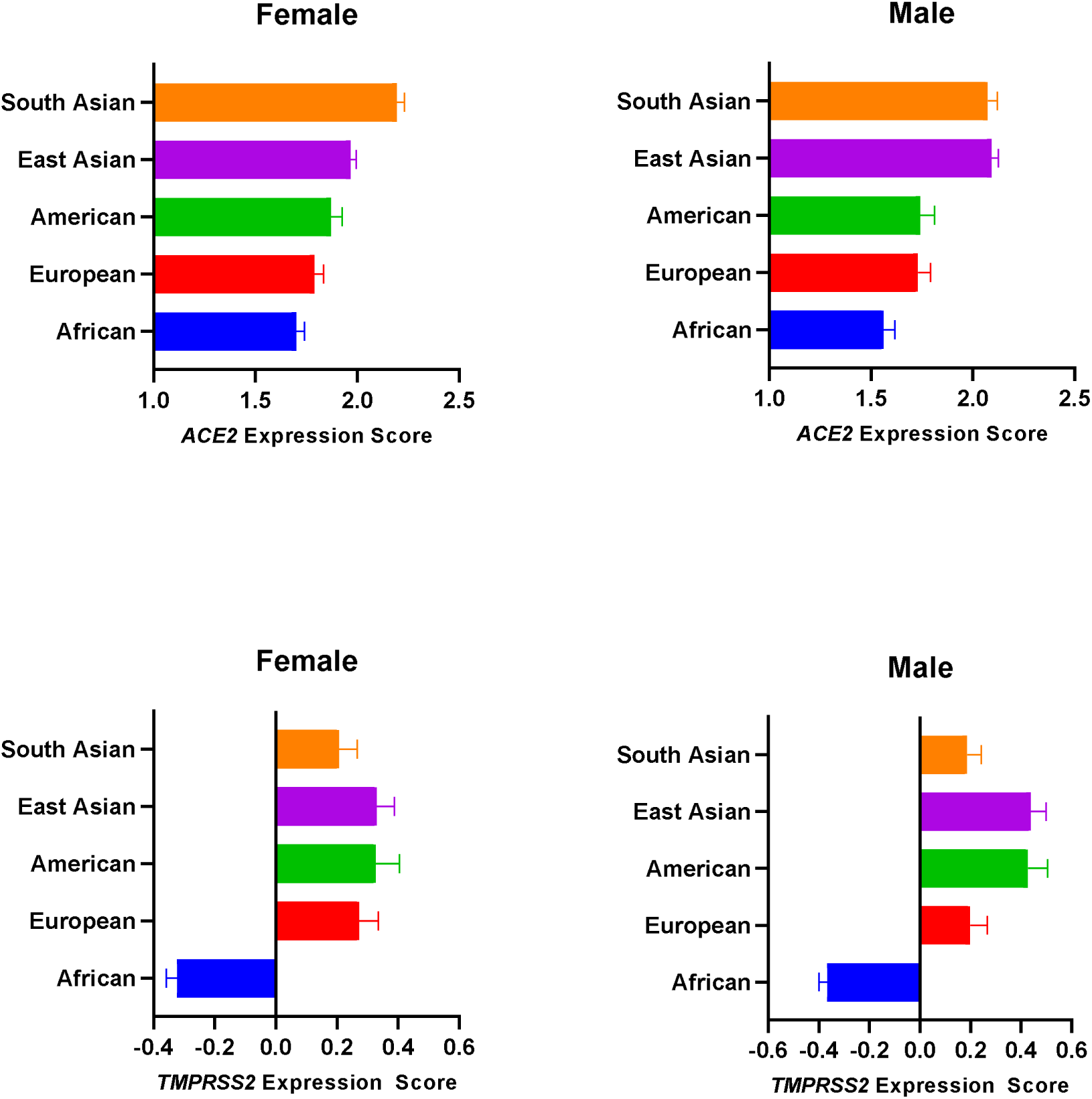
Differences in the cumulative effect of genetic polymorphisms on the expression of SARS-CoV-2 entry mediators *ACE2* and *TMPRSS2* between populations. **(A)** Genetic risk expression score of *ACE2* in female individuals. ANOVA *P* <0.0001. Adjusted *P* values using Tukey’s multiple comparisons test: African vs. European, *P*= 0.47; African vs. American, *P*= 0.043; African vs. East Asian, *P* <0.0001; African vs. South Asian, *P* <0.0001; European vs. American, *P*= 0.71; European vs. East Asian, *P*= 0.0175; European vs. South Asian, *P* <0.0001; American vs. East Asian, *P*= 0.56; American vs. South Asian, *P* <0.0001; East Asian vs. South Asian, *P*= 0.0017. **(B)** Genetic risk expression score of *ACE2* in male individuals. ANOVA *P* <0.0001. Adjusted *P* values using Tukey’s multiple comparisons test: African vs. European, *P*= 0.14; African vs. American, *P*= 0.17; African vs. East Asian, *P* <0.0001; African vs. South Asian, *P* <0.0001; European vs. American, *P*= 1; European vs. East Asian, *P* <0.0001; European vs. South Asian, *P* <0.0001; American vs. East Asian, *P*= 0.0004; American vs. South Asian, *P*= 0.0009; East Asian vs. South Asian, *P*= 1. **(C)** Genetic risk expression score of *TMPRSS2* in female individuals. ANOVA *P* <0.0001. Adjusted *P* values using Tukey’s multiple comparisons test: African vs. European, *P* <0.0001; African vs. American, *P*< 0.0001; African vs. East Asian, *P*< 0.0001; African vs. South Asian, *P* <0.0001; European vs. American, *P*= 0.97; European vs. East Asian, *P*= 0.95; European vs. South Asian, *P=* 0.93; American vs. East Asian, *P*= 1; American vs. South Asian, *P*= 0.66; East Asian vs. South Asian, *P*= 0.55. **(D)** Genetic risk expression score of *TMPRSS2* in male individuals. ANOVA *P* <0.0001. Adjusted *P* values using Tukey’s multiple comparisons test: African vs. European, *P* <0.0001; African vs. American, *P* <0.0001; African vs. East Asian, *P* <0.0001; African vs. South Asian, *P* <0.0001; European vs. American, *P*= 0.084; European vs. East Asian, *P*= 0.03; European vs. South Asian, *P=* 1; American vs. East Asian, *P*= 1; American vs. South Asian, *P*= 0.053; East Asian vs. South Asian, *P*= 0.015.

Similarly, significant differences in GRES for *TMPRSS2* were observed in both female and male groups across populations (ANOVA *P* <0.0001). East Asian populations had the highest values for genetic determinants of *TMPRSS2* expression, and Africans showed genetic predisposition for the lowest *TMPRSS2* expression levels across populations (**Figure 1**).

As mentioned earlier, *ACE2* is located on the X chromosome (and not on the pseudoautosomal region). Therefore, female individuals will have two copies of the gene while males will only have one. Normally, X-chromosome genes are subject to random X-chromosome inactivation, silencing one gene copy in females to keep gene expression balance between females and males. However, a number of X-chromosome genes, including *ACE2*, are known to escape X-chromosome inactivation.^8^ We did not observe differences in genetically determined *ACE2* expression between male and female individuals. Previous reports showed higher expression of *ACE2* in male compared to female tissues, and that this difference in expression was predominantly attributed to non-genetic factors, consistent with our findings.^8^ No difference between male and female individuals for *TMPRSS2* genetic risk expression scores were observed in our study.

These data suggest that genetic determinants of *ACE2* and *TMPRSS2* expression might play a role in variability in transmission and severity of SARS-CoV-2 between populations. African populations showed a genetic predisposition for lower expression levels of both *ACE2* and *TMPRSS2*, which are vital for SARS-CoV-2 entry into host cells. These data might help explain lower numbers of reported COVID-19 in Africa, although the involvement of other non-genetic factors cannot be excluded. Developing assessments that include host genetics might help evaluate disease transmissibility, presentation, and prognosis of COVID-19 in individual patients.

## Notes

Conflict of interest: The authors have declared that no conflict of interest exists

